# An amino-terminal point mutation increases EAAT2 anion currents without affecting glutamate transport rates

**DOI:** 10.1101/2020.04.08.031997

**Authors:** Bettina M I Mertens, Daniel Kortzak, Arne Franzen, Christoph Fahlke

## Abstract

Excitatory amino acid transporters (EAATs) are prototypic dual function proteins that function as coupled glutamate/Na^+^/H^+^/K^+^ transporters and as anion-selective channels. Both transport functions are intimately intertwined at the structural level: secondary active glutamate transport is based on elevator-like movements of the mobile transport domain across the membrane, and the lateral movement of this domain results in anion channel opening. This particular anion channel gating mechanism predicts the existence of mutant transporters with changed anion channel properties, but without alteration in glutamate transport. We here report that the L46P mutation in the human EAAT2 transporter fulfils this prediction. L46 is a pore-forming residue of the EAAT2 anion channels at the cytoplasmic entrance into the ion conduction pathway. In whole-cell patch clamp recordings, we observed larger macroscopic anion current amplitudes for L46P than for WT EAAT2. Moreover, changes in selectivity made gluconate permeant in L46P EAAT2. Non-stationary noise analysis revealed unchanged unitary current amplitudes in mutant EAAT2 anion channels. Rapid L-glutamate application under forward transport conditions demonstrated that L46P does not reduce the transport rate of individual transporters. Since individual transport rates and unitary anion current amplitudes are unaffected, absolute open probabilities of EAAT2 anion channels were quantified as ratio of anion currents by glutamate uptake currents. We found up to sevenfold increased absolute open probability of L46P EAAT2 anion channels. Our results reveal an important determinant of the diameter of EAAT2 anion pore and demonstrate the existence of anion channel gating processes outside the EAAT uptake cycle.

---

After release from presynaptic nerve terminals glutamate is quickly removed from the synaptic cleft by a family of glutamate transporters, the excitatory amino acid transporters (EAATs) (1-3). EAATs are not only secondary active glutamate transporters, but also function as anion-selective channels (4-6). EAAT anion channels regulate neuronal excitability and synaptic transmission (7,8) as well as intracellular [Cl^-^]_int_ in glial cells (9). The importance of EAAT anion channels for normal cell function is emphasized by naturally occurring mutations that cause episodic ataxia and epilepsy and alter EAAT anion currents (10-13).

Glutamate transport is based on the translocation of the transport domain that encompasses binding sites for all transporter substrates. After association of glutamate this domain moves across the membrane via a piston-like movement and then releases substrates to the intracellular membrane side. This so-called elevator mechanism was first described for the prokaryotic glutamate transporter Glt_Ph_ (14,15) and later shown to also apply to an increasing number of other secondary active transporters (16-18). For EAAT/Glt_Ph_, it ensures strict stoichiometric coupling of glutamate to Na^+^, K^+^ and H^+^ transport by permitting translocation only for certain ligation states of the transporter (19-22). Elevator-like transport is also the basis of the anion channel function of this class of glutamate transporters. In atomistic molecular dynamics simulations, the lateral movement of the transport domain and subsequent water entry in the cleft between transport and trimerization domain generates a selective anion conduction pathway (23). This novel conformation correctly predicts experimentally determined selectivities among anions and unitary anion currents and accounts for all published mutagenesis results on EAAT anion channels (23-27).

This mechanism of EAAT/Glt_Ph_ anion channel opening permits the evolutionary optimization of transport rates without changes in anion currents and alterations of anion channel opening without modification in transport rates. It thus accounts for the existence of specialized glutamate transporters and low capacity transporters with predominant anion channel function (4,5,28). To further test the predictions of this model, we searched for point mutations that modify anion channel open probabilities without altering glutamate transport rate. In the course of these experiments, we identified one mutation, L46P, which substantially increased macroscopic anion current amplitudes of EAAT2. We did not observe changes in the voltage and substrate dependence of mutant anion currents, suggesting that the glutamate transport cycle remained unaffected by this mutation. We here performed a detailed investigation of the consequences of this mutation on glutamate transport and anion conduction of EAAT2.

## Results

### L46P increases macroscopic EAAT2 anion currents

Fig. 1*A* depicts an alignment of amino-terminal sequences with mammalian SLC1 transporters and prokaryotic homologues. Bacterial transporters usually exhibit shorter amino-terminal domains, and leucine 46 is not conserved among the family. At present, no three-dimensional structure exists for EAAT2, and we therefore mapped the homologous residue, leucine 54, to outward- (29) and inward-facing conformation (30,31) of ASCT2, the only SLC1 transporter with known structures in both conformations (Fig. 1*B*). Leucine 54 is close to the amino-terminal end of the trimerization domain and does not undergo movements during the isomerization from outward-to inward-facing conformations. The open channel conformation has so far only been described for Glt_Ph_ (23). the homologous valine 12 is a pore-lining residue in Glt_Ph_, which projects its side chain into the water-filled anion permeation pathway (23) (Fig. 1D).

**Figure 1.**
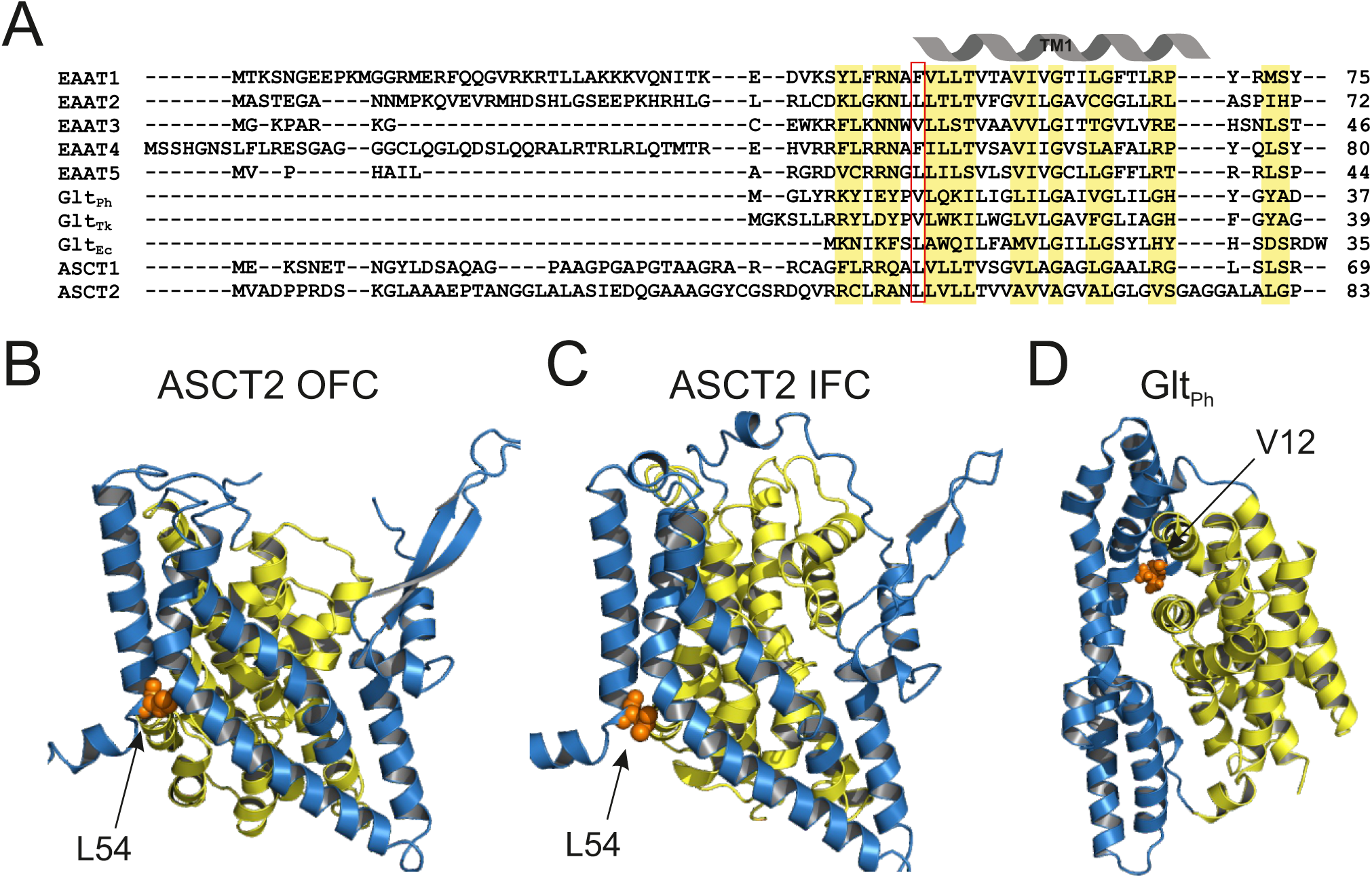
L46 is a pore-forming residue and part of the trimerization domain. *A*, alignment of the N-terminus of EAATs, ASCTs and prokaryotic homologs. Highly conserved regions are marked in yellow. *B*-*D*, position of the L46-homologous residue L54 in the ASCT2 topology model in the outward (29), inward (30,31) and of V12 in the Glt_Ph_ open channel conformation (23) (blue: trimerization domain; yellow: transport domain).

Fig. 2, *A* and *B* depict representative recordings from cells transiently transfected with WT (Fig. 2*A*) or L46P EAAT2 (Fig. 2*B*). In these experiments, cells were dialyzed with KNO_3_-based internal solution and then perfused either with NaNO_3_-based external solutions without or with 1.0 mM L-glutamate, or with an external solution, in which NaNO_3_ was completely substituted with KNO_3_. EAAT2 transports glutamate in the presence of external Na^+^ and L-glutamate (19-21). The use of NO_3_^-^ that is more permeant than Cl^-^ (32) increases EAAT2 anion currents to levels greatly exceeding uptake currents (5,33) (Fig. 2, *C* and *D*). Almost complete block upon application of TBOA, which prevents opening of EAAT anion channels by blocking inward movement of the transport domain (34), demonstrates that WT and mutant EAAT2 anion currents are significantly larger than background currents (Fig. 2, *C* and *D*) and can thus be directly measured without subtraction procedures (33,35).

**Figure 2.**
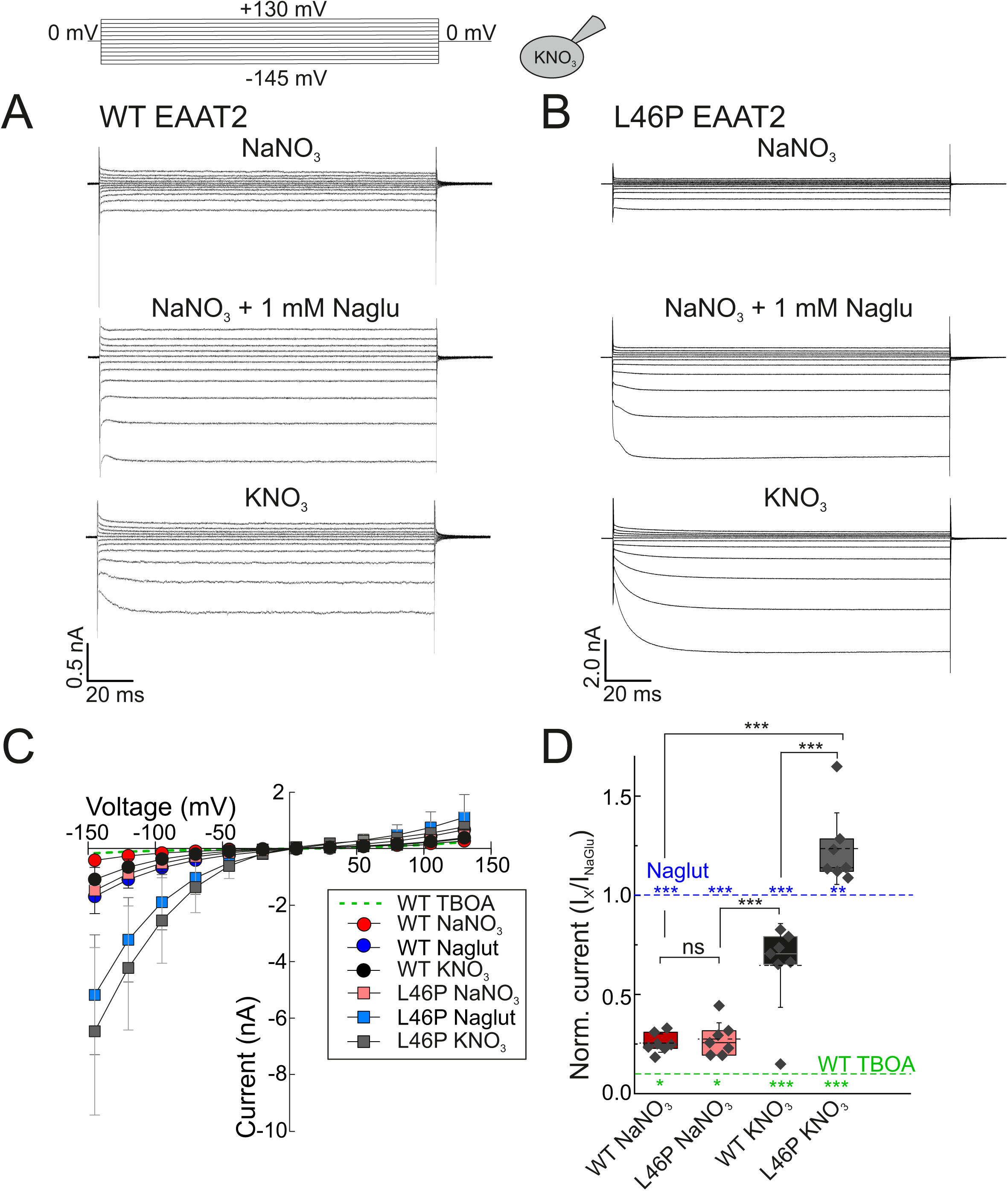
L46P increases EAAT2 anion currents under a variety of ionic conditions. *A* and *B*, representative current recordings from HEK293T cells expressing WT (*A*) or L46P (*B*) EAAT2. Cells were intracellular dialyzed with a KNO_3_-based solution and perfused with NaNO_3_-based solution without or with glutamate, with TBOA or with KNO_3_-based solutions. *C*, voltage-dependence of mean steady-state currents from cells expressing WT or L46P EAAT2. The background current (dotted, green line) was obtained from cells expressing WT EAAT2 perfused with TBOA. *D*, normalized current amplitudes at −145 mV from cells expressing WT or L46P EAAT2. All error bars represent 95% C.I. with n=7.

WT as well as L46P EAAT2 anion currents are small and without appreciable time and voltage dependence in NaNO_3_-based external solutions lacking glutamate (Fig. 2). Application of L-glutamate increases anion currents for WT and mutant transporters (Fig. 2, *C* and *D*). EAAT2 also mediates large anion current amplitudes under conditions, in which extracellular Na^+^ is completely substituted by K^+^ (36,37). WT and mutant currents displayed similar time, voltage and substrate dependences. However, L46P EAAT2 anion currents are much larger than WT currents. Moreover, whereas WT EAAT2 anion current amplitudes were highest in external Na^+^ and L-glutamate, we observed maximum mutant current under KNO_3_-based external solutions (Fig. 2*D*).

### L46P neither affects expression nor subcellular distribution of EAAT2

To test whether increased macroscopic L46P EAAT2 currents arise from changes in subcellular localization or in expression levels we employed confocal imaging and protein biochemistry (Fig. S1). WT and L46P EAAT2 display predominant surface membrane insertion in confocal images from HEK 293T cells (Fig. S1, *A* and *B*). Fig. S1*C* depicts fluorescent scans of SDS PAGE from membrane preparation of transfected HEK 293T cells, indicating core- and complex glycosylated WT and L46P EAAT2 (38). Individual bands were quantified with the ImageJ software and revealed similar expression levels (Fig. S1*C*). To exclude the possibility that changes in the number of transfected cells compensate for altered expression levels, we also determined transfection rates by manually counting transfected and untransfected cells. We did not observe differences between WT and mutant transporters (Fig. S1*D*) and conclude that L46P leaves expression levels and subcellular distributions of EAAT2 unaffected.

### L46P leaves unitary current amplitudes of EAAT2 anion channels unaltered

Since expression levels of WT and mutant transporters are similar, the observed differences in L46P EAAT2 macroscopic current amplitude might be due to increased mutant anion channel conductance or open probability. We employed non-stationary noise analysis to determine unitary current amplitudes of WT and L46P EAAT2 anion channels (39). To perform such experiments in cells with comparable WT and mutant current amplitudes, we generated inducible stable cell lines for WT as well as for L46P EAAT2 (38).

Fig. 3, *A* and *B* show representative time courses of mean macroscopic currents and corresponding current variances for WT (*A*) or L46P (*B*) EAAT2 elicited by repetitive voltage steps to −140 mV in symmetrical NaNO_3_ and 0.5 mM external L-glutamate. Fig. 3*C* provides pooled noise analysis data from 6 cells expressing WT (red circles) and 17 cells expressing L46P EAAT2 (blue circles). We plotted ratios of current variances by mean current amplitudes against normalized mean current amplitudes at various time points. A fit of the resulting relationship with a linear function (Eq. 2) provides the single-channel current amplitudes as y-axis intercept (40). Bootstrap analysis was used to simulate the error-generating process of data sampling from a larger population and to get an approximation of the true error of fitting parameters. 50,000 bootstrap samples (each containing 300 WT or 850 mutant EAAT2 data points with replacements) were randomly generated from the original data set (39,41) and individually fitted with linear functions (Fig. 3*C*). Single-channel current amplitudes were determined for each bootstrap sample from linear fits, providing single channel amplitudes of 26 ± 2 fA (mean ± 95% C.I.; n = 6) for WT and 29 ± 2 fA (mean ± 95% C.I.; n = 17) for mutant EAAT2 (p > 0.05) (Fig. 3*D*). We conclude that L46P does not modify the unitary current amplitude of EAAT2 anion channels, in agreement with earlier data on EAAT4 carrying a glutamate substitution at the homologous F55 residue (23).

**Figure 3.**
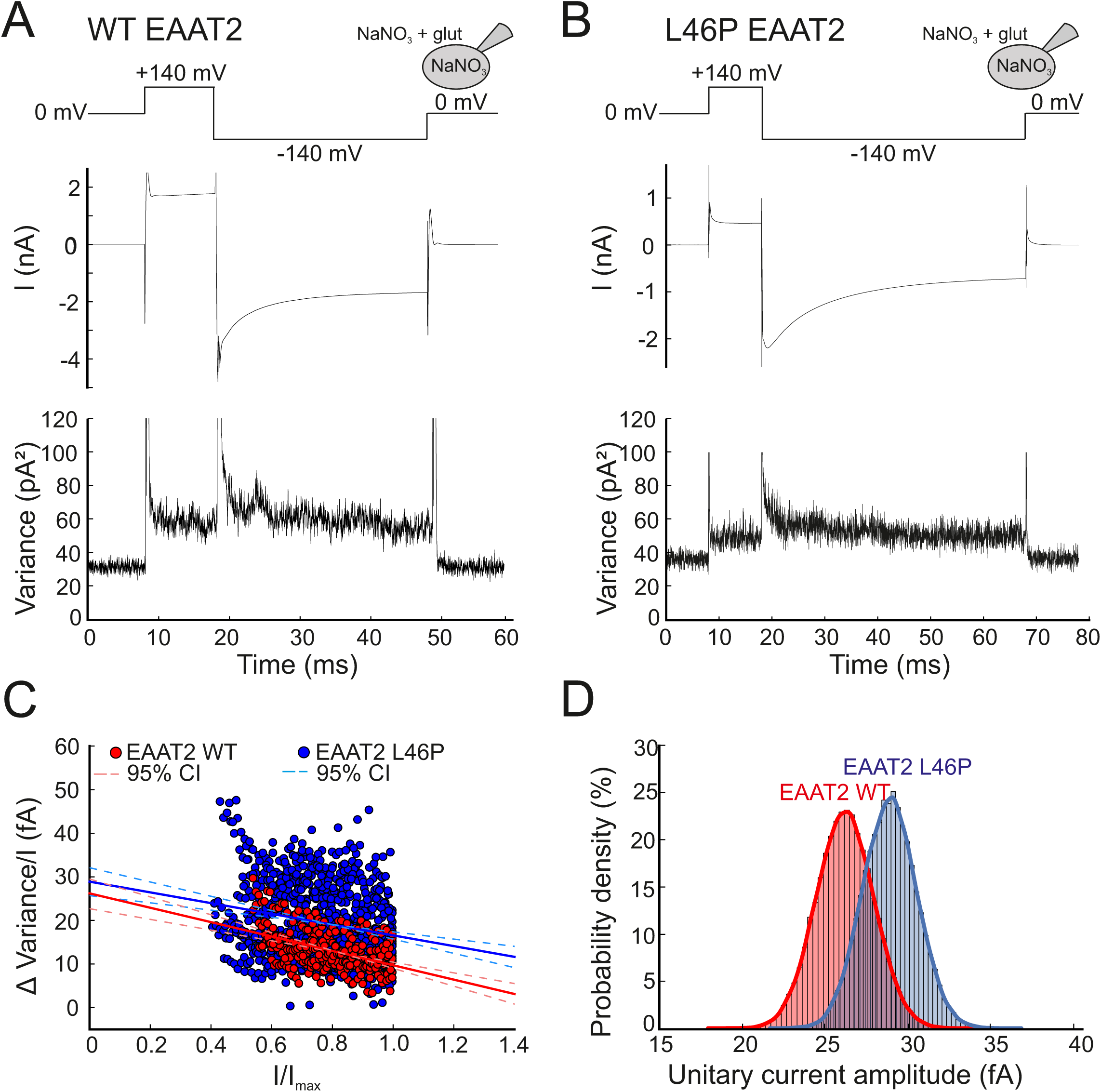
L46P leaves unitary current amplitudes unaffected. *A* and *B*, voltage protocol and time courses of representative current amplitudes and current variances from HEK293T cells transiently transfected with WT (*A*) or Flp-In T-Rex 293 cells stably expressing L46P (*B*) EAAT2 measured in symmetrical NaNO_3_ based solutions. *C*, pooled noise analysis data from 6 cells expressing WT (red circles) and 17 cells expressing L46P EAAT2 (blue circles). For comparison between different cells, current variance by mean current amplitude ratios are plotted against normalized current amplitudes. Solid lines represent fits with linear functions and dashed lines give the 95% confidence interval of the fit determined by bootstrap regression analysis. *D*, distribution of estimated unitary current amplitudes for WT and L46P EAAT2 derived from 50,000 bootstrap samples of the original data. Histograms have been normalized such that the integral over the range is 1 and fitted with a Gaussian function (WT EAAT2: µ=26.19, σ=1.73; L46P EAAT2: µ=28.9, σ=1.64).

### L46P affects pore dimensions of EAAT2 anion channels

EAAT anion channels conduct a variety of anions, they select, however, against large anions by size (23,32,33). The minimum pore size of the Glt_Ph_ anion channel is around 5-6 Å (23), only slightly smaller than the minimum size required for gluconate permeation (6.9 Å, (42)). To test for potential changes in selectivity of L46P EAAT2 anion channels, we recorded currents in cells dialyzed with NaNO_3_ and perfused with solutions containing mixtures of NaNO_3_ and Nagluconate or of NaNO_3_ and CholineNO_3_ (Fig. 4). Outward currents were only negligible in cells expressing WT transporters perfused with NO_3_^-^-free external solutions (Fig. 4, *A* and *C*), in full agreement with a negligible gluconate^-^ permeability. In contrast, we observed significant outward currents for L46P EAAT2 (Fig. 4*D*). Gluconate^-^ permeability of L46P EAAT2 suggests that the mutations widens the selectivity filter and permits passing of anions that are usually too large to permeate.

**FIGURE 4.**
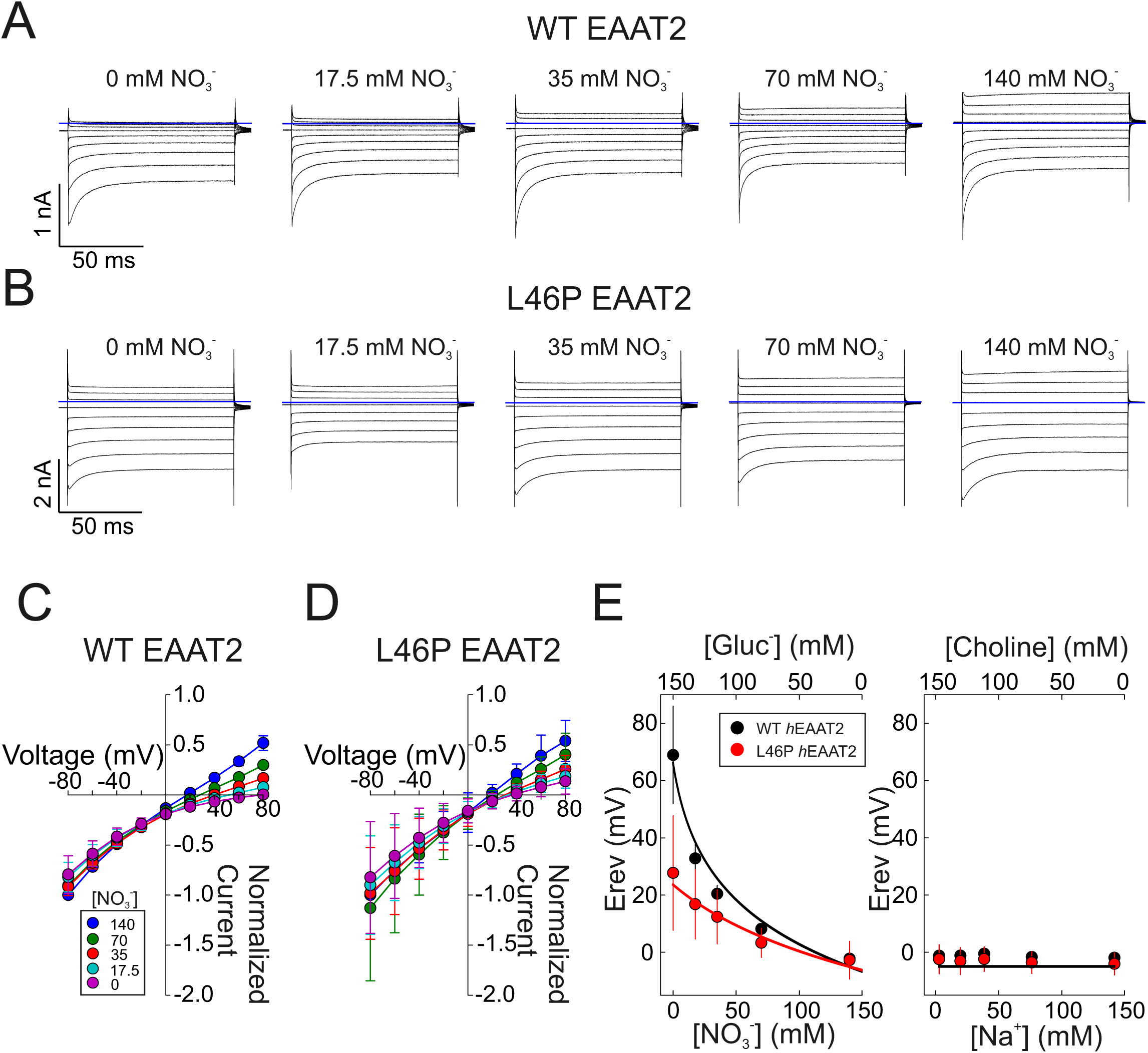
L46P EAAT2 anion channels are permeable to gluconate. *A,B*, representative anion current recordings from single HEK293T cells, which either express WT (*A*) or L46P (*B*) EAAT2, subsequentially perfused with 1 mM glutamate and different external [NO_3_^-^] (substituted with gluconate) and internally dialyzed with a NaNO_3_ based solution. *C,D*, normalized current-voltage curves from cells expressing WT (*C*) or L46P (*D*) EAAT2 at various external [NO_3_^-^]. *E*, changes in WT or L46P EAAT2 anion current reversal potentials upon variation of external [NO_3_^-^] or [Na^+^]. All error bars indicate mean ± S.D..

Plotting current reversal potentials versus external [gluconate^-^] or external [Na^+^] (Fig. 4*E*) demonstrate that WT and mutant transporters differed significantly in their relative gluconate permeability, whereas neither WT nor L46P EAAT2 are measurably permeant to Na^+^. Permeability ratios obtained from the Goldman-Hodgkin-Katz equation provided P_Gluc_/P_NO3_ values of 0.31 ± 0.06 (mean ± S.D. from 50,000 bootstrap samples) for L46P, as compared to 0.07 ± 0.01 (mean ± S.D. from 50,000 bootstrap samples) in WT transporters. The obtained value of the relative gluconate permeability of WT EAAT2 anion channels appears to contradict the absent gluconate inward currents in Fig. 4*A*. The very low macroscopic current amplitudes at very positive potentials for WT EAAT2 likely result in an overestimation of P_gluc_ because even very leak conductances can significantly contribute to the observed reversal potentials.

Surprisingly, no L46P EAAT2 currents were observed in symmetric gluconate (Fig. 6*A*). We thus postulate that NO_3_^-^ binding may induce a transient widening of the L46P EAAT2 selectivity filter, which allows for subsequent gluconate^-^ permeation.

**Figure 5.**
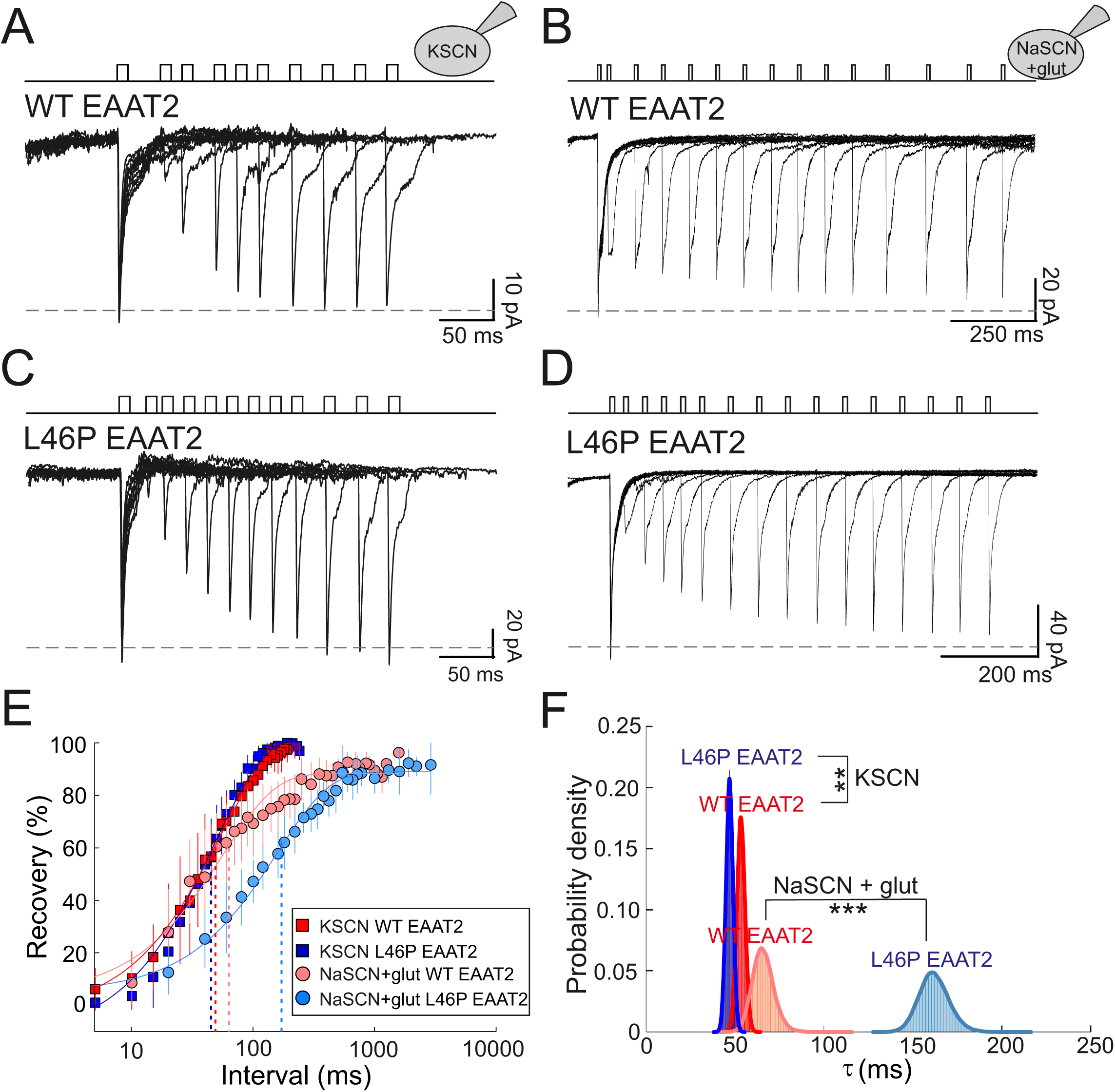
L46P leaves EAAT2 forward transport unaffected. *A-D, r*epresentative rates of recovery from depression of transporter currents from patches containing WT (*A,B*) or L46P (*C,D*) EAAT2. Experiments were performed either with KSCN- (*A,C*) or NaSCN/L-glutamate-based (*B,D*) cytoplasmic solutions. *E*, mean ratios of second response by first response peak current amplitudes plotted against the interval between pulses for 115 mM [KSCN]_in_ from patches containing WT (red squares, τ=53.3 ± 2.3 ms, *n* = 10) or mutant EAAT2 (blue squares, τ=46.9 ± 2.0 ms, n = 8), or for 115 mM NaSCN/10 mM L-glutamate-based solutions from patches containing WT (pink circle. τ=65.9 ± 6.0 ms, *n* = 8) or mutant EAAT2 (light blue circle, τ=161.9 ± 8,4 ms, n = 6). Error bars indicate mean ± S.D.. *G*, Probability density of transport rates obtained with 50,000 bootstrap samples randomly selected from the original data set.

**Figure 6.**
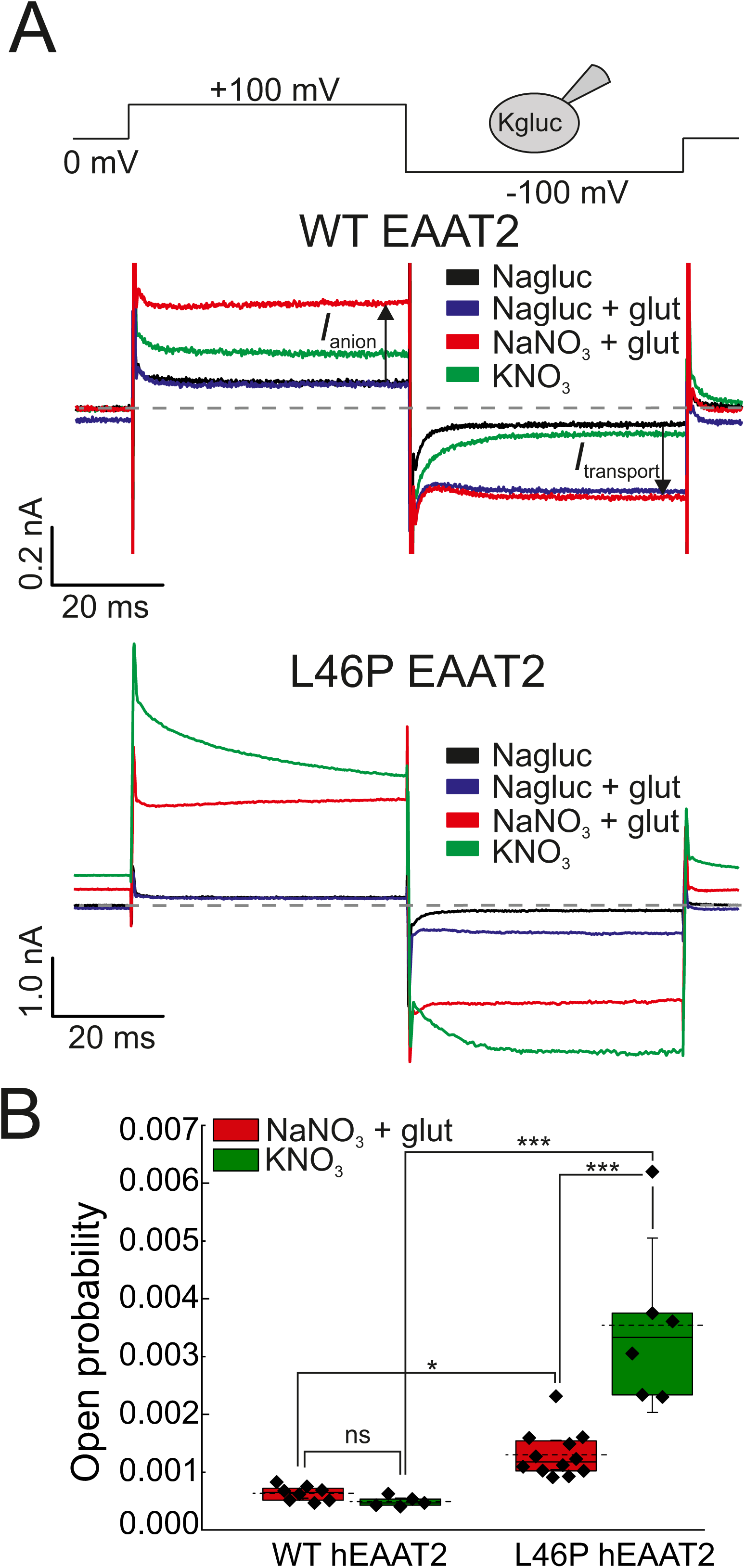
L46P increases absolute open probabilities of EAAT2 anion channels. *A* and *B*, representative current responses of HEK293T cells expressing WT (*A*) or L46P (*B*) EAAT2 to consecutive voltage steps to +100 mV and −100 mV. Cells were dialyzed with a Kgluconate-based pipette solution and consecutively perfused with Nagluconate-based solution with or without L-glutamate, with a NaNO_3_-based solution containing 0.5 mM L-glutamate, and finally with a KNO_3_-based solution. *C*, estimated anion channel open probabilities for WT and mutant EAAT2 anion channels at Na^+^- or K^+^-based external solutions. Data are shown as mean ± 95% C.I..

### WT and L46P exhibit similar unitary transport rates

WT and L46P EAAT2 neither differ in unitary anion currents nor in expression levels, making enhanced absolute open probabilities of mutant anion channels a likely reason for the large anion currents in cells expressing L46P EAAT2. Recent work has demonstrated that non-stationary noise analysis cannot accurately determine absolute open probabilities of EAAT anion channels (11,43). We therefore decided to estimate the fraction of time, in which EAAT transporters assume the open anion channel conformation, using consecutive measurements of coupled transport and anion currents at the same cell (43). Since this analysis requires not only knowledge about unitary anion current amplitudes, but also of individual transport rates we measured recovery rates of transport current depression using fast substrate application on excised outside-out patches to measure these values (44,45).

Under conditions permitting forward transport, i.e. intracellular K^+^ and extracellular Na^+^, fast application of L-glutamate initiates transport cycles and generates a transient capacitive current followed by a steady-state transport current (Fig. 5). The amplitudes of both current components are proportional to the number of transporters accessible to external glutamate. In a double pulse protocol with two subsequent glutamate applications, the amplitude of the second electrical signal depends on the time period between the two pulses. If the second pulse is applied immediately after the first pulse, the majority of transporters will assume intermediate positions of the transport cycle and only few transporters will be accessible to external glutamate. With increasing duration between two subsequent glutamate applications, the current amplitude will recover with a time course that is determined by the average time required for a complete transport cycle (44,45).

Fig. 5 depicts representative current responses to subsequent 10 ms application of 10 mM L-glutamate at increasing intervals for WT (Fig. 5, *A* and *B*) and L46P EAAT2 (Fig. 5, *C* and *D*). With K^+^-based pipette solutions, the recovery was fitted with a single exponential function with time constants of 53.3 ± 2.3 ms (mean ± S.D. from 50,000 bootstrap samples; n=10) for WT and slightly faster values (46.9 ± 2.0 ms (mean ± S.D. from 50,000 bootstrap samples; n =8), p < 0.005) for L46P EAAT2 (Fig. 5, *E* and *F*). We conclude that L46P causes only slight changes in rate-limiting steps of forward glutamate transport.

We next performed comparable experiments under homoexchange conditions, with intracellular solutions containing Na^+^ and L-glutamate (Fig. 5, *B* and *D*). Under these conditions, full transport cycles are impossible and recovery rates are determined by re-translocation and unbinding of L-glutamate. Under this condition L46P decelerated the time course of recovery (65.9 ± 6.0 ms (mean ± S.D. from 50,000 bootstrap samples; n=8) for WT and 161.9 ± 8.4 ms (mean ± S.D. from 50,000 bootstrap samples; n=6) for L46P EAAT2) (Fig. 5, *E* and *F*). Forward transport and homoexchange transport cycles differ in the re-translocation step from inward to outward-facing conformations. Since recovery time constants are significantly longer under homoexchange conditions than under forward transport, the re-translocation step must be the rate-limiting transitions in both cases. Thus, L46P decelerates L-glutamate-bound re-translocation and/or substrate dissociation from the outward-facing conformation of EAAT2 transporters.

### Absolute open probabilities of L46P EAAT2 anion channels are significantly increased

To compare EAAT2 glutamate uptake and anion currents at individual cells, we studied whole-cell currents in cells intracellularly dialyzed with a Kgluconate-based pipette upon subsequent perfusion with Nagluconate- and then with NaNO_3_- or KNO_3_-based external solutions (43) (Fig. 6). Electrogenic glutamate transport can be measured in isolation as glutamate-induced EAAT2 currents when permeant anions are completely substituted with gluconate (5). Under these conditions, we observed comparable current amplitudes for WT and L46P EAAT2 at −100 mV (Fig. 6), in full agreement with unaltered expression and subcellular distribution (Fig. S1) and only slight differences in individual transport rates of mutant transporters (Fig. 5). However, application of L-glutamate supplemented NaNO_3_-based external solution revealed dramatic difference in WT and mutant anion current amplitudes at positive potentials (+100 mV). Due to the pronounced voltage dependence of glutamate uptake currents (5) these currents are predominantly conducted by EAAT2 anion channels. Subsequent change to KNO_3_-based external solutions resulted in current amplitudes that are only slightly smaller than for Na^+^/glutamate in WT and much larger in L46P EAAT2 (Fig. 6, *A* and *B*).

To obtain the number of transporter subunits in the studied cells, we divided the transport current amplitude at −100 mV by WT or L46P transport rates (Fig. 6*C*). We next applied this number together with the unitary anion current amplitude obtained in Fig. 3 to calculate absolute open probabilities for WT and mutant EAAT2 (43). We determined an absolute open probability of 0.06 % ± 0.01 % (mean ± 95% C.I.; n = 9) for WT and 0.13% ± 0.03% (mean ± 95% C.I.; n = 12) for L46P EAAT2 at +100 mV with external NaNO_3_ and L-glutamate. L46P thus causes twofold larger open probability for EAAT2 anion currents than for WT under forward transport conditions and a sevenfold increase under K^+^-exchange conditions (0.05% ± 0.01% (mean ± 95% C.I.; n = 5) for WT and 0.35% ± 0.15% (mean ± 95% C.I.; n = 6) for L46P EAAT2 at +100 mV) with external KNO_3_ (Fig. 5*C*). Since L46P EAAT2 does not reach a steady state current under K^+^-exchange conditions, an exponential function was fitted and the instantaneous current 1 ms after pulse application was used to calculate the open probability.

External application of NaNO_3_ or KNO_3_ to cells expressing L46P EAAT2 did not only increase outward currents at positive potentials, but also inward currents at negative voltages (Fig. 6*B*). Such currents, which were absent in cells expressing WT EAAT2, are due to the gluconate permeability through L46P EAAT2 anion channel in the presence of NO_3_^-^ (Fig. 4).

## Discussion

Computational and experimental data support the notion that EAATs/Glts form anion-selective conduction pathways at the interface between trimerization and transport domain via lateral movement of the transport domain in intermediate conformations (23,26,27). Progression of the transport cycle results in closure of the anion channel (23), and efficient glutamate transport is only possible at the cost of low anion channel open probability. Functional analysis of *SLC1A3* mutations associated with episodic ataxia type 6 revealed that excessive EAAT anion currents might perturb brain development and function (10-13). The interdependency between glutamate transport and anion channel activity together with the pathological consequences of increased EAAT anion channel activity demonstrate the evolutionary need for a strict regulation of anion channel gating.

We identified one mutation in EAAT2 that substantially increases macroscopic anion current amplitudes, without discernible effects in its voltage and substrate dependence and studied the functional consequences of this L46P mutation on glutamate transport and unitary properties of EAAT2 anion channels. We demonstrate that L46P leaves protein expression and subcellular distribution of EAAT2 unaffected (Fig. S1), but augments anion currents (Fig. 2). The increased anion current amplitudes are not due to changes in unitary current amplitudes (Fig. 3), but rather caused by an increased probability of occupying open anion channel conformations (Fig. 6). Moreover, L46P causes pronounced effects on EAAT2 anion channel selectivity, resulting in significant permeability of gluconate^-^ (Fig. 4). L46P leaves unitary current amplitudes unaffected (Fig. 3), but increases gluconate permeability (Fig. 4). This finding is in agreement with the notion that anions do not directly interact with L46 (23), but rather modify the open anion channel conformation, resulting in larger pore diameters.

The dual function of EAATs as transporter and as anion channel and the resulting complex gating mechanism underlying EAAT anion channel opening makes the determination of absolute open probabilities difficult. For many ion channels, non-stationary noise-analysis accurately quantifies unitary current amplitudes together with the number of channels in the evaluated cell/membrane patch. However, this method does not provide reliable numbers for EAATs. Noise analysis on EAAT anion currents provided rather large open probabilities (33,35), with values well above 0.5. Recently, a disease-causing point mutation was reported that decreases the number of transporters in the surface membrane, but increases macroscopic anion current amplitudes more than fivefold (11) at unaltered unitary current amplitudes. This observation demonstrates that absolute open probabilities of EAAT anion channels are not correctly determined using noise analysis. A possible reason for this inaccuracy has been already described for the anion-proton exchanger ClC-4 that also assumes anion channel modes in addition to functioning as secondary active transporter (40). Very fast transitions in combined transport/anion channel cycles cause semi-equilibria between transporters with open and with closed channels. Hence, only transporters, which undergo rapid transitions between open and closed channel, will be counted by noise analysis, resulting in underestimation of the number of transporters and overestimation of absolute open probabilities (40).

We used an alternative approach that is based on the comparison of glutamate transport currents and anion currents (43). We first determined transport rates of WT and L46P EAAT2 (Fig. 5), and then used these transport rates to calculate the numbers of transporters in individual cells from measured glutamate uptake currents (Fig. 6). Dividing macroscopic anion current amplitudes by the corresponding transporter number and the unitary current amplitude (Fig. 3) provides absolute open probabilities (Fig. 6). This analysis reveals that WT EAAT2 spend only 0.06 % of the total time in the anion conducting state under ionic conditions that permit forward transport, and an absolute open probability of 0.05 % in the K^+^-exchange mode. These values are very low, and are fully consistent with EAAT2 being an effective glutamate transporter with rather low associated anion current (5). L46P increases this value to 0.13% in the forward mode, and to 0.35% in the K^+^ exchange mode.

Our analysis of L46P EAAT2 illustrates that changes in anion channel open probability can occur without concomitant alterations in glutamate uptake. We determined individual transport rates by measuring recovery rates of transporter current depression in the forward transport mode (Fig. 4) and observed closely similar values for WT and L46P EAAT2. L46P thus does not cause major changes in forward transport rates. However, L46P EAAT2 anion currents – either in absolute values (Fig. 2) or after normalization to transport rates (Fig. 6) - are much larger. L46P thus modifies transitions between intermediate transport conformation and open channel conformation (23).

Our estimates of glutamate transport rates resemble data with native neuronal EAATs in hippocampal Purkinje neurons (46), but are smaller than published values on heterologously expressed (47) or native (44) EAAT2. At present, we do not know the origin for these differences. They might be due to small variation in the composition of the internal and external solutions. Otis and Kavanaugh use higher internal [K^+^] and apply TEA to the cytoplasmic membrane, rather than to the external solution as we did in our experiments (47). There are earlier reports on variation of recovery rates depending in cell systems and experimental approach (45). L46P has a slightly more pronounced effect on anion current amplitudes in the K^+^ exchange mode than under glutamate forward transport conditions (Figs. 1 and 6). This might be due to a K^+^ dependence in anion channel opening. Alternatively, they might indicate subtle alterations of certain transport transitions. L46P has only minor effects on recovery rate of depression of the transporter current under forward transport conditions (Fig. 5), indicating similar glutamate transport rates for WT and mutant transporters. However, under Na^+^-glutamate exchange rates, recovery time constants of L46P EAAT2 are more than twofold larger than for WT (Fig. 5). These differences suggest that L46P additionally affects Na^+^/glutamate-bound re-translocation and/or release. Neither in an inward-facing ASCT2 structure nor in the EAAT1 structure, L46P is directly interacting with the transport domain. However, it is in close proximity of the gate to the substrate binding sites (HP2) so that local conformational changes could translate in different translocation or glutamate release kinetics.

Earlier work comparing glutamate transport and anion current amplitudes permitted the distinction of EAAT isoforms that mainly operate as glutamate uptake carriers (EAAT1, EAAT2, EAAT3) (5) and others, EAAT4 and EAAT5, that exhibit low glutamate transport rates and predominantly function as glutamate-gated anion channels (4,48). Published data on EAAT1 and EAAT3 depict smaller anion than uptake currents (5), indicating very small anion channel open probabilities also for these two isoforms. Such experiments have not been possible for EAAT4 or EAAT5 because of the low glutamate transport rates of these two isoforms (28,49). However, the recent comparison of a mouse EAAT2 splice variant, GLT-1c, and mouse EAAT5 in the same expression system with identical experimental approaches for both isoforms (39) revealed similar macroscopic anion current amplitudes at comparable expression levels and subcellular distribution. This finding argues against major differences in absolute open probability between high and low capacity glutamate transporters. EAAT4 and EAAT5 have been shown to exhibit very small transport rates, suggesting that the main difference between specialized EAAT transporters and EAAT anion channels is the transport rate rather than the anion channel open probability (28,49). However, the small open probability of EAAT anion channels permits adjustment of open probabilities without change in transport rates.

In summary, we here present a point mutation that modifies EAAT2 anion channel open probabilities without significant alteration of transport rates. The point mutation exchanges a pore-forming leucine residue located at the cytoplasmic entrance to the predicted EAAT/Glt_Ph_ anion conduction pathway. We interpret the effects on anion channel open probabilities by proposing that L46P affects the energy difference between intermediate transport conformation and open anion channels. The increased pore diameter of L46P EAAT2 resulting in measurable gluconate^-^ permeability suggests distinct positions of the transport domain in open anion channel conformation of WT and L46P EAAT2. Our findings provide additional experimental evidence for the recently proposed EAAT/Glt_Ph_ anion conduction mechanism (23).

## Experimental procedures

### Heterologous expression of WT and mutant EAAT2

WT (kindly provided by Dr. M. Hediger, University of Bern, Switzerland) and mutant human EAAT2 were expressed as mYFP fusion protein as described previously (36). mYFP-EAAT2 fusion proteins have been intensively tested and been shown to be functionally indistinguishable from EAAT2 lacking the fluorescent protein (36). The L46P point mutation was introduced using PCR-based strategies, and the resulting pRcCMV-mYFP L46P EAAT2 was verified by restriction analysis and DNA sequencing. Two independent recombinants from the same transformation were examined and shown to exhibit indistinguishable functional properties. Transient transfection of HEK293T cells using the Ca_3_(PO_4_)_2_ technique was performed as previously described (50,51). Transfection rates were determined by manually counting transfected and untransfected cells on confocal images (2386 cells (3 transfections) for WT and 2211 cells (3 transfections) for L46P). Since macroscopic anion currents often exceeded 10 nA in cells transiently expressing L46P EAAT2, stable inducible cell lines expressing WT or mutant EAAT2 were generated by selection of Flp-In T-Rex 293 cells (Invitrogen) transfected with pcDNA5/FRT/TO-EAAT2 and used either with or without induction with tetracycline (38,50,52).

### Electrophysiology

Standard whole-cell or outside-out patch clamp recordings were performed using an HEKA EPC10 amplifier (HEKA Electronics, Lamprecht, Germany) as described (53). Borosilicate pipettes were pulled with resistances of 1.5-3.0 MΩ. To reduce voltage errors, we routinely compensated more than 80% of the series resistance by an analog procedure and excluded recordings with more than 10 nA maximum anion currents from the analysis. In experiments with outside-out patches, pipettes were covered with dental wax to reduce their capacitance. Currents were filtered at 2.9 kHz (−3dB) and digitized with a sampling rate of 50 kHz. Cells and patches were clamped to 0 mV for at least 4 s between test sweeps.

To record anion currents (Figs. 2, 3 and 4) the pipette solution contained (in mM) 115 Na/KNO_3,_ 2 MgCl_2_, 5 EGTA, 10 HEPES, pH 7.4. The control extracellular bath solution contained (in mM) 140 NaNO_3_ ± 1.0 L-glutamate. Under potassium-bound homoexchange conditions extracellular NaNO_3_ was equimolar replaced by KNO_3_. For rapid application experiments pipette solution contained (in mM) 115 NaSCN + 10 L-glutamate under glutamate-bound homoexchange conditions or 115 KSCN under transport conditions, and the bath solution 140 NaNO_3_, 4 KCl, 2 CaCl_2_, 1 MgCl_2_, 5 HEPES, pH 7.4. To measure open probabilities under transport conditions or under homoexchange conditions an internal solution, in which Na/KNO_3_ were equimolarly substituted with K-gluconate was used. To block macroscopic current 0.1 mM DL-TBOA was added to a NaNO_3_-based solution.

### Fast substrate application

For fast application of transport substrates, a piezo-driven system with a dual-channel theta glass (45,54) tubing was used (Fig. 4) (MXPZT-300, Siskiyou, Grants Pass, OR, USA). At the end of each experiment we removed the cell from the patch pipette and measured 20-80 % rise times of our system by fast application of the used solution to the open pipette (54). We obtained values between 1.3 and 3.0 ms, with a mean value of 2.2 ± 0.4 ms (mean ± C.I.; n=30).

### Noise analysis

We used non-stationary noise analysis to determine single-channel current amplitudes of WT and L46P EAAT2 anion channels (Fig. 3) (39). Current variances were calculated from current differences in subsequent records during 300 subsequent voltage jumps (39) and subtracted the background noise measured at 0 mV. The 10% of the sweeps with the highest time-averaged variances were removed.

Ion channels usually generate a Lorentzian-type of noise. Since open and closed states of an ion channel are binomially distributed, the amplitude and the time dependence of the current variance (*σ*^*2*^) can be calculated by

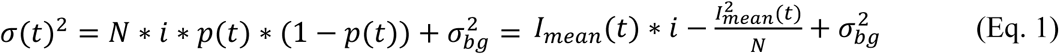

or after linear transformation

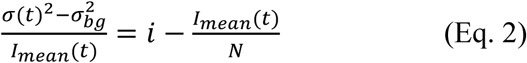

with *i* being the single-channel current amplitude, *p* the absolute open probability, *N* the total number of channels in the membrane, *I*_*mean*_ the mean macroscopic current amplitude and *σ*^*2*^_*bg*_ the voltage-independent background noise. The y-axis intercept from a linear regression of Eq. 2 to the data points thus provides the unitary current amplitude, and the slope of the linear regression *(-N*^*-1*^) the number of channels.

We normalized mean currents (*I*_*mean*_) to the maximum current amplitudes (*I*_*max*_) to account for differences in transporter expression and pooled normalized data from all cells expressing WT or mutant EAAT2 into single plots. We plotted ratios of current variances by non-normalized current amplitudes versus normalized current amplitudes for all examined cells (Fig. 3 *C*). The regression error was assessed by bootstrap sampling simulations (41). For this, 50,000 bootstrap samples were resampled by randomly selecting data from the original data sets with replacement. We determined single-channel amplitudes for all bootstrap samples and the distribution of these values was plotted in a histogram for visual inspection (Fig. 3 *D*). The reported errors of the single-channel currents are the confidence intervals of this parameter of the regression of all bootstrap samples.

### Confocal imaging

Images were acquired 24-36 h after transfection with a Leica TCS SP5 II inverted microscope (Mannheim, Germany) using a 63x oil immersion objective from living cells in PBS containing Ca^2+^ and Mg^2+^ at room temperature (22-24°C) (55). mYFP (monomeric yellow fluorescence proteins) fluorophores were excited with a 488-nm Argon laser. Confocal images were assembled for publications in ImageJ (56).

### Biochemical Analysis

Membrane proteins were solubilized with 0.4% DDM after membrane isolation from transfected HEK293T using hypotonic and hypertonic solutions (57). Total protein concentrations were determined using the BCA assay, and cleared lysates were then denatured for 15 min at room temperature in SDS sample buffer and electrophoresed in parallel with fluorescent mass markers (Dual Color, Bio-Rad) on 10% SDS polyacrylamide gradient gels. YFP-tagged proteins were visualized by scanning the wet PAGE gels with a fluorescence scanner (Typhoon, GE Healthcare, München, Germany). Individual bands were quantified with ImageJ (53).

### Data analysis

Data analysis was performed using a combination of FitMaster (HEKA), self-written programs in Phython (Continuum Analytics), Origin (OriginLab), SigmaPlot (Systat Software) and Excel (Microsoft) software. All data is presented as mean ± 95% confidence interval or standatd deviation (S.D.) as indicated. For fast-application and noise analysis, S.D. was assessed by bootstrap sampling simulations with 50,000 bootstrap samples, randomly selected from the original data set. In boxplot diagrams, boxes represent 25.75 percentiles, whiskers indicate 95% confidence intervals, straight lines show the median value and dotted lines the mean value. Mean values were compared using two-way ANOVA with Holm-Sidak *post hoc* test. Data are significant different with p⩽0.05 (*), p⩽0.01 (**) or p⩽0.001 (***).

## Acknowledgments

These studies were supported by the Deutsche Forschungsgemeinschaft to Ch.F. (FA 301/12-1) as part of the Research Unit FOR 2518, DynIon; project P4.

---

**Supplemental Figure S1.**
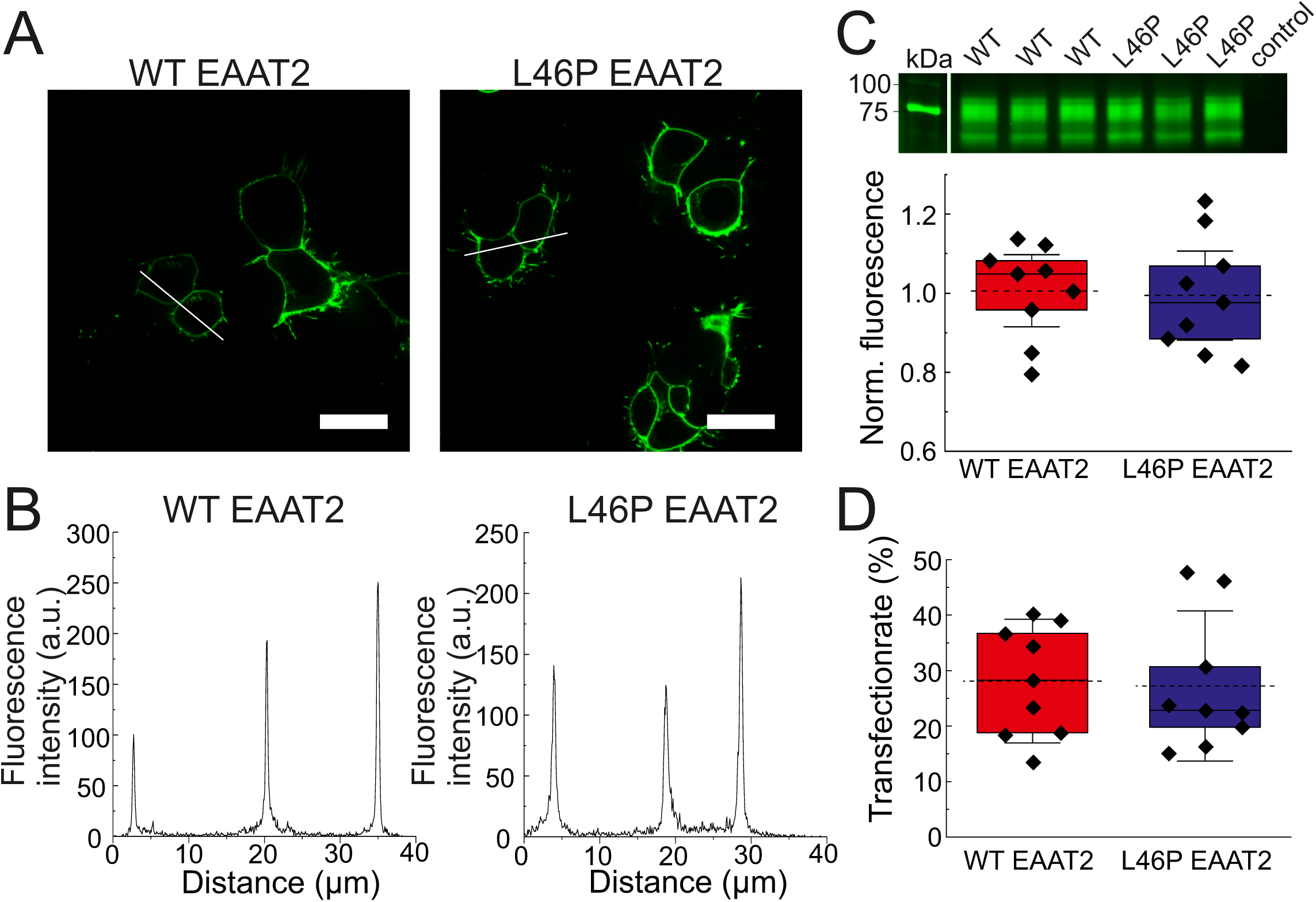
L46P leaves EAAT2 expression and subcellular distribution unaffected. *A* and *B*, confocal images (*A*) and corresponding intensity profiles (*B*) of HEK 293T cells transiently transfected with WT or L46P EAAT2. Scale bar 25µm. *C*, representative fluorescence scan of a SDS-PAGE gel and mean fluorescences from whole cell membrane fractions from HEK 293T cells expressing WT or L46P EAAT2. Data are normalized to the mean fluorescence of the respective gel. *D*, transfection rates of cells expressing WT or L46P EAAT2. Data are presented as mean ± 95% C.I..

